# Epitope mapping of SARS-CoV-2 RBDs by hydroxyl radical protein footprinting reveals the importance of including negative antibody controls

**DOI:** 10.1101/2023.10.27.564378

**Authors:** Daniel Nyberg Larsen, Jakub Zbigniew Kaczmarek, Yaseelan Palarasah, Jonas Heilskov Graversen, Peter Højrup

**Affiliations:** Department of Biochemistry and Molecular Biology, University of Southern Denmark, Campusvej 55, DK-5230 Odense, Denmark; Ovodan Biotech A/S, Havnegade 36, DK-5000 Odense, Denmark; Department of Cancer and Inflammation, Institute of Molecular Medicine, Faculty of Health and Medical Sciences, University of Southern Denmark, Odense, Denmark

**Keywords:** RBD Alpha, RBD Delta, Mass spectrometry, Fenton chemistry, Covid-19

## Abstract

Understanding protein-protein interaction is essential when designing drugs or investigating biological processes. A variety of techniques can be employed in order to map the regions on proteins that are involved in binding eg., CryoEM, X-ray spectroscopy, linear epitope mapping, or mass spectrometry-based methods. The most commonly utilized mass spectrometry-based techniques are cross-linking and hydrogen-deuterium exchange (HDX). An alternative technique for identifying residues on the three-dimensional structure of proteins, that are involved in binding, can be hydroxyl radical protein footprinting (HRPF). However, this method is currently hampered by high initial cost and complex experimental setup. Here we set out to present a generally applicable method using Fenton chemistry for mapping of epitopes in a standard mass spectrometry laboratory. Furthermore, the described method illustrates the importance of controls on several levels when performing mass spectrometry-based epitope mapping. In particular, the inclusion of a negative antibody control has not previously been widely utilized in epitope mapping by HRPF analysis. In order to limit the number of false positives, we further introduced quantification by TMT labelling, thereby allowing for direct comparison between sample conditions and biological triplicates. Lastly, up to six technical replicates were incorporated in the experimental setup in order to achieve increased depth of the final analysis.

Both binding and opening of regions on receptor-binding domain (RBD) from SARS-CoV-2 Spike Protein, Alpha, and Delta variants, were observed. The negative control antibody experiment combined with the high overlap between biological triplicates resulted in the exclusion of 40% of the significantly changed regions, including both binding and opening regions. The final identified binding region was mapped to a three-dimensional structure and agrees with the literature for neutralizing antibodies towards SARS-CoV-2 Spike Protein.

The presented method is straightforward to implement for the analysis of HRPF in a generic MS-based laboratory. The high reliability of the data was achieved by increasing the number of technical and biological replicates combined with negative antibody controls.

## 1 Introduction

In December 2019 a novel coronavirus (SARS-CoV-2) crossed species barriers to infect humans and has effectively been transmitted from person to person [1]. Infection with SARS-CoV-2 causes the coronavirus disease 19 (COVID-19) with influenza-like symptoms, ranging from mild disease to server lung injury and multi-organ failure, potentially leading to death for elderly patients with other co-morbidities [2,3].

SARS-CoV-2 infects the host by a densely glycosylated spike (S) protein [2]. The S protein is a trimeric class I fusion protein that exists in two different conformations and undergoes a substantial structural rearrangement to fuse the viral membrane with the host-cell membrane [2]. The fusion is triggered when a subunit of the S protein (S1) binds to the host-cell receptor Angiotensin converting enzyme 2 (ACE2). The region responsible for spike protein binding to ACE2 is the Receptor-Binding Domain (RBD). The binding destabilizes the prefusion trimer, resulting in a post-fusion conformation. To engage the host the viral receptor undergoes hinge-like conformational movements that transiently hide or expose regions used during receptor binding, these two states are referred to as the “down” and “up” conformation, where “down” corresponds to the receptor-inaccessible state and “up” corresponds to the receptor-accessible state [2]. The region used during receptor binding is found to be closer to the central cavity of the trimer, compared to the closely related beta-coronaviruses [2].

As the underlying mechanism of the SARS-CoV-2 infection, the interaction between RBD of the spike protein and the ACE2 receptor has automatically been targeted for therapeutic antibody development. A wide variety of neutralizing monoclonal antibodies have been generated from hybridoma [4–6] or immunized organisms using recombinant technologies [7,8] and have shown promising effects in inhibitory studies and a few have already been approved for treatment by the FDA [9,10].

Detailed knowledge of the specific interactions of pairs of antibodies and antigens, the epitope and paratope, respectively, are important during selection of antibodies for further development of therapeutic or prognostic use [11]. Furthermore, precise knowledge strengthens the IP protection of a given mAb. For the development of therapeutic mAbs detailed knowledge of the epitope is a prerequisite for conducting clinical trials [12] and to increase the comparability between animal and human studies when surrogate mAbs are to be used due to lacking cross-reactivity between species, both for safety and for proof-of-concept. A wide arsenal of techniques for the characterization of protein-protein binding interfaces is available [13,14]. Often these technologies complement each other and are used in combination to ensure highly confident results. However, the majority of the methods require prior assumptions of the structure of the interaction and are very labor intensive, making the methods unsuitable for early-stage Ab screening and selection. The methods used today are X-ray crystallography, Cryo electron microscopy, mutational scanning, hydrogen / deuterium exchange (HDX-MS) [14–17], chemical cross-linking (CX-MS) [18–22], and chemical foot-printing [23–27]. These methods all require several sample handlings steps, which might alter the native structure of the protein in question.

Currently, the gold standard for Ab epitope mapping is X-ray crystallography. This is cumbersome work, with a very limited throughput. Thus, novel and more efficient methods are being sought. One such approach is based on hydrogen/deuterium-exchange in combination with MS. This has to some degree reduced the time needed to map epitopes, however, the method is relatively complicated and requires detailed data analysis. From the introduction of Fast Photochemical Oxidation of Proteins (FPOP) from Michael Gross’s laboratory [25,28], the Hydroxyl Radical Protein Footprinting (HRPF) has been utilized in epitope mapping. After introduction into the field several other alternatives have been implemented and especially with FOX systems [29], PLIMB [30], and X-ray-activated hydroxyl radicals [27] showing promising results. However, the main problem with these systems is the need for specialized equipment, which is expensive to acquire and maintain. Fenton chemistry [31,32], that was first introduced in 1876 and 1934, use the reaction of Fe2+ with H_2_O_2_ to generate hydroxyl radicals. The hydroxyl radicals are one of the most powerful, nonselective, irreversible, and fast reacting radicals, which interact with both organic and inorganic compounds. The main reason that the field has not yet implemented the method is the slow reaction time of seconds to minutes, which can cause the protein to unfold or degrade upon oxidation. This is unlike FPOP, that has a reaction time in the millisecond time scale.

The use of controls within HRPF is essential to distinguish background from introduced oxidation, which is achieved by utilizing a sample that has not been subjected to oxidations [28,33]. Incorporation of biological replicates ensures that small variations do not create false positive results. Technical replicates should be utilized to minimize the variations between biological replicates. However, technical replicates are not generally reported [23,29,30,34,35] for this methodology. Furthermore, the analysis of antibody binding epitopes by HRPF does not generally include non-binding antibodies (negative antibodies). The negative antibody control could be useful in removing false positive results (binding regions/paratope) when the interaction between the nonsense antibody and screened antigen occurs. The use of negative antibodies is very common in regular antibody experimental methodologies, as unspecific binding of nonsense antibodies is generally observed [36,37].

Commercially available instrumentation for epitope mapping is currently available but is not commonly used due to the high costs. In order to get the epitope mapping into the general mass spectrometric laboratory the overall costs must be decreased, or the methods coupled to existing infrastructure. This could be pipetting robots which in addition to higher throughput, increase reliability and reproducibility. Within this work, we have combined two different methods with different analysis methodologies, using surface plasmon resonance (SPR) and HRPF. From SPR two antibodies were selected to verify the binding and non-binding antibody, thereby allowing for the use of a true negative antibody control. HRPF was used as a method based on Fenton chemistry integrated into a semi-automized workflow on a pipetting robot. The combination of SPR and a semi-automated MS workflow allows to utilize the speed of SPR and the precision of HRPF. Biological triplicate analysis was performed for both the negative antibody control and the binding antibody. Furthermore, all samples were run in technical sextuplicate to elucidate the technical variance. Stable isotope labeling in the form of TMT labelling was used as a quantification method, which is in contrast to the generally used label-free quantification [23,28,38].

The focus of the present study was epitope mapping of a mouse monoclonal antibody to the receptor-binding domain of two SARS-CoV-2 spike protein variants (Wild-Type and variant B.1.617.2). The experimental setup utilizing multiple controls allowed for validation of false positive binding on several levels, while still being easily implementable in the general MS laboratory. The primary equipment needed to implement this method in addition to a mass spectrometer is a pipetting robot e.g., Opentrons OT-2.

## 2 Material and methods

### 2.1 Antigen production

The gene synthesized for expression of the SARS-CoV-2 Spike RBD 319-541 WT (Wild-Type) variant has been based on the sequence GISAID: EPI_ISL_402119 (referred to as “Alpha” in this article). Variant B.1.617.2 (commonly and further in this article referred to as ‘’Delta’’) has been based on the sequence EPI_ISL_2029113. Both constructs were synthesized commercially and cloned into pcDNA3.4-TOPO plasmid (Life Technologies B.V. Europe) that included a Tyr-Pho signal peptide and an N-terminal hexahistidine (6xHis) tag. For both variants the expression was carried out in 25 mL culture using the Expi293F expression system (#A14635; ThermoFisher Scientific) according to the manufacturer’s instructions. Proteins harvested after 6 days post transfection were purified from the supernatant on a Ni-NTA Superflow column (#30430, Qiagen). Immediately after the elution, protein fractions were buffer exchanged into PBS pH 7.4 using a HiPrep 26/10 desalting column (#GE17-5087-01, Cytiva) and stored at -20°C. The purity and size of the protein has been evaluated by Sodium Dodecyl Sulfate Polyacrylamide gel electrophoresis (SDS-PAGE) using RunBlue (#ab270467 Abcam) 4-12% Bis-Tris polyacrylamide gel. Electrophoretic separation has been performed using Easy Power 500 (Invitrogen) in a non-reduced environment for 45 minutes at 200V at 110mA. The gel was stained with Coomassie Simply Blue Safe Stain (#LC6060 Invitrogen) and developed according to the manufacturer’s instructions. Both SARS-CoV-2 Spike RBD 319-541 variants WT (#OBA0101, Ovodan Biotech) and variant B.1.617.2 (#OBA0105, Ovodan Biotech) have been donated by Ovodan Biotech A/S, Denmark (data sheets available in supplementary data).

### 2.2 Antibody

Monoclonal antibodies (mAbs) specific for the receptor-binding domain of two SARS-CoV-2 variants were generated by immunizing NMRI mice with recombinant RBD of SARS-CoV-2 (RBD, His Tag) (Sinobiological, number 40592-V08B). The mice were immunized twice subcutaneously, at an interval of 14 days, with 20 μg of the mixed RBD with GERBU adjuvants (GERBU Biotechnik GmbH, Heidelberg, Germany). Fourteen days after the last injection and three days prior to fusion, the mice were immunized intravenously with recombinant RBD, and spleen cells were harvested and fused with myeloma cells, following the principles of Köhler and Milstein [39]. Supernatants from hybridomas were initially screened using Maxisorb ELISA plates (ThermoFisher Scientific, Roskilde, Denmark) coated with 0.5 µg/mL purified recombinant RBD followed by at least three rounds of cloning by limited dilution, ensuring clonality of the final hybridoma. Single clones were then grown in culture flasks in RPMI-1640 (Lonza, BioWhittaker, Basel, Switzerland) supplemented with sodium pyruvate (Gibco™, Waltham, MA, United States) and gentamicin sulfate (Biowest, Nuaillé, France) containing 10% fetal bovine serum (Biowest). Previously developed and characterized antibody against peptidylarginine deiminase 4 (PAD4) was used as negative control.

### 2.3 Surface plasmon resonance analysis

Surface plasmon resonance (SPR) analysis was performed on a Biacore T200 instrument (Cytiva) and analyzed using Biacore T200 Evaluation Software (Cytiva). RBD variants were immobilized on a CM5 sensor chip using EDC/NHS chemistry at 10 µg/ml in a 10 mM acetic acid pH 5 buffer according to the manufacturer’s instructions.

Binding studies were performed at a 10 µl/minute flow in the following running buffer: 140 mM NaCl, 10 mM Hepes, 0.1 Tween 20 pH 7.4 and the flow cells were regenerated with two injections of 20 µl, 100 mM H_3_PO_4_ between runs. All experiments were performed at least as triplicates of technical duplicates.

### 2.4 Samples

Biological triplicate samples were all prepared in a 96-well plate (Axygen, Corning incorporated), where they were lyophilized. Two experimental setups were run right after the other, the Control experiment (containing RBD Alpha and non-binding anti-PAD4 antibodies) and the Sample experiment (containing RBD Alpha and anti-RBD 20-14-5). Three sample types were prepared for both experiments: antibody, RBD Alpha or Delta, and a 1 to 1.5 molar ratio RBD and antibody. Each starting well contained 15 µg of protein.

### 2.5 Oxidative labelling

The oxidative labelling was performed on the Opentrons OT2 pipetting robot (Opentrons Labworks, Inc.), where the thermocycler module were used for cooling the sample. The robot was partially sealed and pressurized with nitrogen gas to decrease the oxygen level within the robot.

The samples were incubated in 32 µl footprinting buffer (FB) (10mM MOPSA/NaOH, 200mM NaCl, 10mM MgCl2) for 15 min. at 25⁰C, followed by aliquoting of the negative control (0 min incubation) sample into a new 96-well plate located in the thermocycler at 4⁰C. Each utilized well in the 96 well plate contained quenching buffer (QB) (0.1 M L-methionine) (10 µl for time zero and 20 µl for 3 and 9 min). Oxidative labelling was then prepared by addition of 6 µl Iron-EDTA (20 mM EDTA, 20 mM ammonium iron (II) sulfate hexahydride, in FB) (buffer mixed 30 min before use) and 6 µl 0.2 M sodium ascorbate. The labelling was then initiated by the addition of 6 µl 10 mM freshly made hydrogen peroxide. All buffers were degassed prior to use, except for QB. After 3 min of incubation, half the volume of each sample was transferred to the microtiter plate in the thermocycler into 20 µl QB. The remainder of the samples were transferred after 9 min of incubation time to the thermocycler. To the 0 min incubation time sample 20 µl of 100 mM HEPES was added.

### 2.6 Sample preparation

The oxidative labelled samples were removed from the robot, and the rest of the preparations were performed by hand. The samples were reduced by the addition of dithiothreitol (DTT) to a final concentration of 20 mM and incubated for 30 min at 57⁰C. This was followed by alkylation of the cysteines by the addition of Iodoacetamide (IAA) to a final concentration of 54 mM and incubated at RT in the dark for 20 min. The alkylation was stopped by the addition of 1 µl 0.1 M DTT.

The digestion was performed in three steps. 2 µl PNGaseF (P0705L, Glycerol-free, New England Biolabs) was added and incubated for one hour at 57⁰C. 2% LysC w/w (125-05061, Lysyl Endopeptidase Mass Spectrometry Grade, Fujifilm Wako Chemicals) was added, and the incubation continued for an hour at 37⁰C, and finally 2% methylated trypsin (methylated in-house) [40] was added and incubated for 10 hours at 37⁰C.

Samples were labelled by Tandem Mass Tag (TMT) 11-plex according to the manufacturer’s instructions, where 10 tags were used. Furthermore, one common channel was used containing 1.5 µg of each sample from both experiments. All 10 tags were mixed in equal ratios for each biological replicate. The resulting 6 tubes containing 3 biological replicates for the Control and Sample experiment were micro-purified in accordance with Rappsilber et al. [41], followed by lyophilization. For each biological replicate 6 purifications were performed resulting in a total of 36 lyophilized samples.

### 2.7 Mass spectrometry

Each sample was resuspended in 5.5 μl 0.1% formic acid prior to mass spectrometric analysis and transferred to a 96 well microtiter plate and loaded into an EASY-nLC^TM^ 1200 system (Thermo Fisher Scientific). The LC system was connected to an Exploris 480 Orbitrap mass spectrometer (Thermo Fisher Scientific). Each biological replicate was run in six technical replicates using a 60 min gradient and the following main MS setting: Full scan: Orbitrap resolution of 120,000, scan range (m/z) of 350-1500, normalized AGC Target (%) of 300, maximum Injection Time Mode was set to Auto. Filters applied were MIPS: Monoisotopic peak determination of Peptide and Relax restrictions was set to True; Intensity: Filter type was Intensity Threshold set to 5.0e3; Data Dependent: Mode was Number of Scans with number of scans set to 10. Scan Event Type 1; Charged states used was 2-7, and undetermined charge states were not included; Dynamic exclusion: Exclude after 1 time, excluded for 60 seconds, mass tolerance was ±10ppm, Isotopes were excluded. Data dependent MS2 settings used were isolation window 0.8 m/z, HCD collision energy of 30%, Orbitrap resolution of 45,000, First Mass of 110 m/z, Standard AGC Target, Maximum Injection Time Mode was Auto, Data type was Centroid.

The samples were run with a BSA standard for every third sample and three BSA standards between every biological replicate. The samples were run as follows: Sample experiment biological replicate 1 (SBRep1), Control experiment biological replicate 1 (CBRep1), SBRep2, CBRep2, SBRep3, and CBRep3.

### 2.8 Data analysis

The resulting 36 raw files from each experiment, 36 from RBD Alpha and 36 from RBD Delta, were searched in Proteome Discoverer 2.5 (Thermo Scientific). The files were recalibrated by quick search followed by several filtering steps. The spectra were filtered by signal to noise level of 1.5 and 10 peaks pr. 100 Da. Three rounds of searches were performed, the first two by the MASCOT search engine and the last with Sequest HT. Identified PSMs with a medium confidence or lower, were searched again for each round. The main search settings were 10 ppm precursor mass tolerance, 0.1 Da fragment mass tolerance, and two missed cleavages allowed. The following modifications were used for the MASCOT searches, for the first: Deamidation (NQ), Oxidation (P, F, D, M) Di-oxidation (M, W), and Carbonyl (S, V); and for the second: Deamidation (NQ), Oxidation (P, H, W, M, R, S), and Carbonyl (S, L, I). The last search by Sequest HT used the following modifications: Deamidation (NQ), Oxidation (D, F, H, K, M, N, P, Q, R, S, T, W, Y), Trioxidation (F, W, Y), and Carbonyl (A, E, I, L, P, Q, R, S, V). Ratios were calculated between all the conditions and the common channel and used for analysis. The resulting ratios for peptide groups were exported as CSV files including amongst other the following data: sequence, modifications in all possible sites, ratios, protein accession number, and molecular weight. Furthermore, the full modification list was exported for RBD Alpha or Delta depending on the experiment.

The data were imported into R studio [2023.03.0+386, Posit PBC] using an R markdown script, where the data were filtered and analyzed. Peptide groups were accepted if they were found in at least two out of three biological replicates and found in a single technical replicate pr. biological replicate. Calculations of p-values using a two-sided Student’s t-test were utilized. Ratios were average across the three biological replicates both for original data and for 3/0 min incubation and 9/0 min incubation, these were used for part of the analysis.

Ratio changes between antigen and antigen:antibody (Mix) were calculated by dividing the normalized abundance for 3 min or 9 min incubation with abundance for 0 min incubation within each sample type (Ag, Ab, or Mix). The values were then divided again between Mix and Ag or Mix and Ab, followed by averaging across the three biological replicates. For non-oxidized peptide values above 1 in the ratio between Mix and Ag or Mix and Ab indicate binding or shielding of the region, whereas values below 1 indicate an opening of the region. For oxidized peptides values below 1 in the ratio between Mix and Ag or Mix and Ab indicate binding or shielding of the region, whereas values above 1 indicate an opening of the region [42].

The mass spectrometry proteomics data have been deposited to the ProteomeXchange Consortium via the PRIDE [43] partner repository with the dataset identifier PXD046423.

## 3 Results

### 3.1 Antigen production

Two variants of the receptor-binding domain of the SARS-CoV-2 spike protein were expressed and purified. The alpha variant (RBD Alpha) and the delta variant (RBD Delta) were expressed and screened for correct location of mutations by mass spectrometry, where 93% coverage was achieved without PNGaseF and 100% coverage with PNGaseF for WT. This showed the presence of glycosylation on either or both potential sites, which are located on the same tryptic peptide. Expressed proteins were additionally tested by ELISA using anti-SARS-CoV-2 polyclonal IgY (OBP1121, Ovodan Biotech) confirming the presence of the protein. The size of the expressed RBD variant proteins has been determined by SDS-PAGE showing the expected size of 37 KDa for double glycosylated RBD aa 319-541. (**Error! Reference source not found.**).

### 3.2 Binding characterization

To characterize the binding between either RBDs to the anti-RBD antibody (20-14-5) and demonstrate the lack of binding to the negative control antibody (PAD4 14-11-3), binding kinetics were performed by SPR on both antigens and antibodies. The mAb 20-14-5 exhibited binding to both RBD Alpha and Delta in the nM range with comparable affinities and kinetics, see Figure 1B. No binding to the control mAb (PAD4 14-11-3) was observed, see Figure 1A.

**Figure 1.**
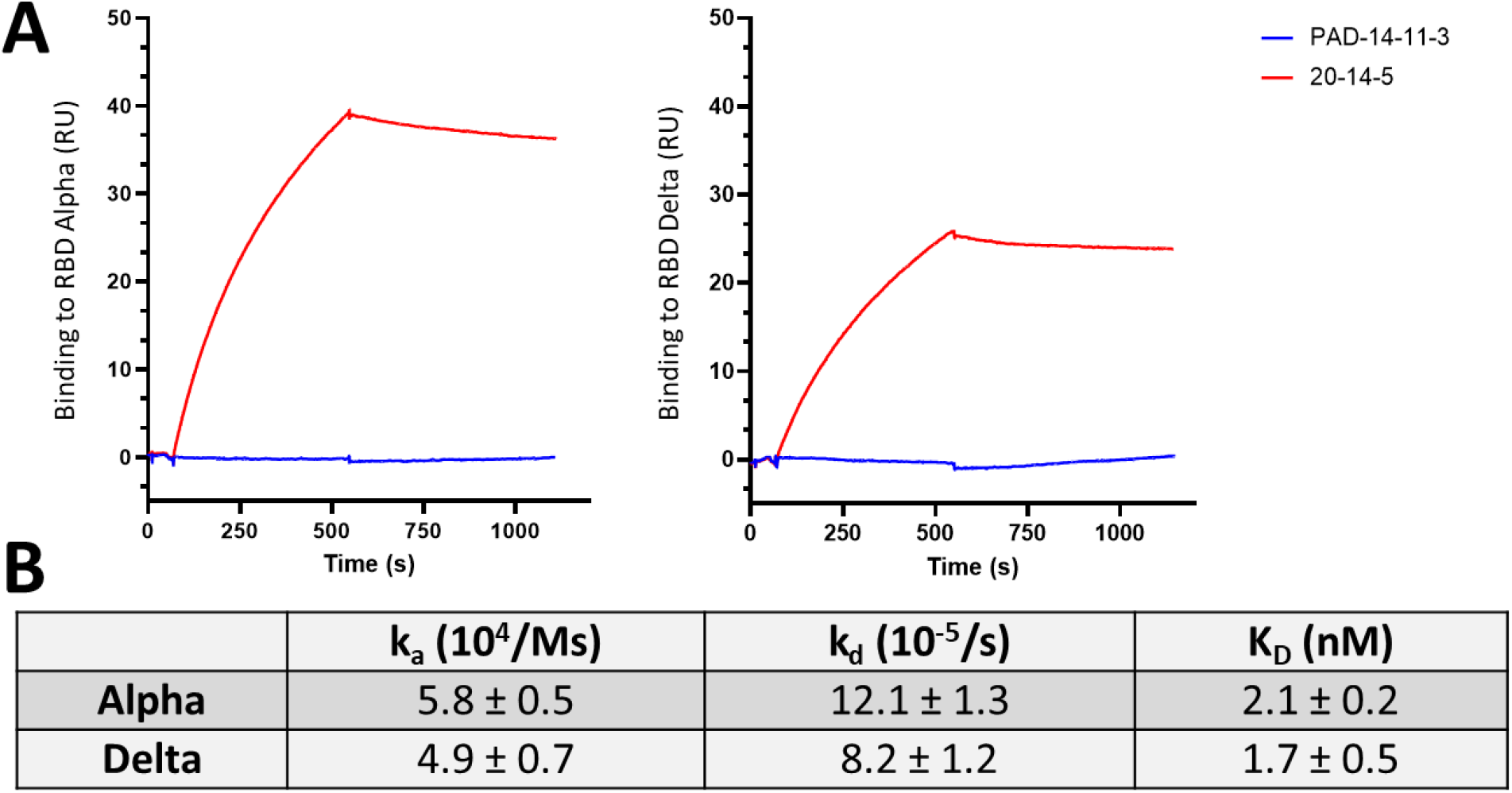
A) SPR analysis of binding of 5 µ/ml 20-14-5 and control mAb (PAD-14-11-3) to RBD Alpha and Delta immobilized at densities of 0.036 and 0.041 pmol/mm2, respectively. B) Binding parameters for the binding of 20-14-5 to RBD Alpha and Delta, using 20-14-5.

### 3.3 Epitope mapping

The data generated from oxidative labelling were processed and tested for reproducibility with regard to biological and technical replicates. Technical replicates were evaluated in Figure 2A and E for RBD Alpha and Delta, respectively. For RBD Alpha the identification across all types of peptide groups shows an increasing trend for each additional technical replicate, where replicate 2 contributes to an increase of 19.5%, and the following 13.9%, 10%, 8.2%, and 3.6% respectively. For RBD Delta the increases in identification from the technical replicate were 25%, 8.7%, 5.3%, 3%, and 7.8%, respectively. The largest increase was observed for the first and second replicates for both RBD Alpha and Delta, whereas less increase was observed for the last three replicates. The full overview of increases by subcategory are presented in Supplementary Figure 1. A continual increase was expected, however, less increase by additional replicates indicates that the depth of analysis is high and that additional replicates result in more identifications.

**Figure 2.**
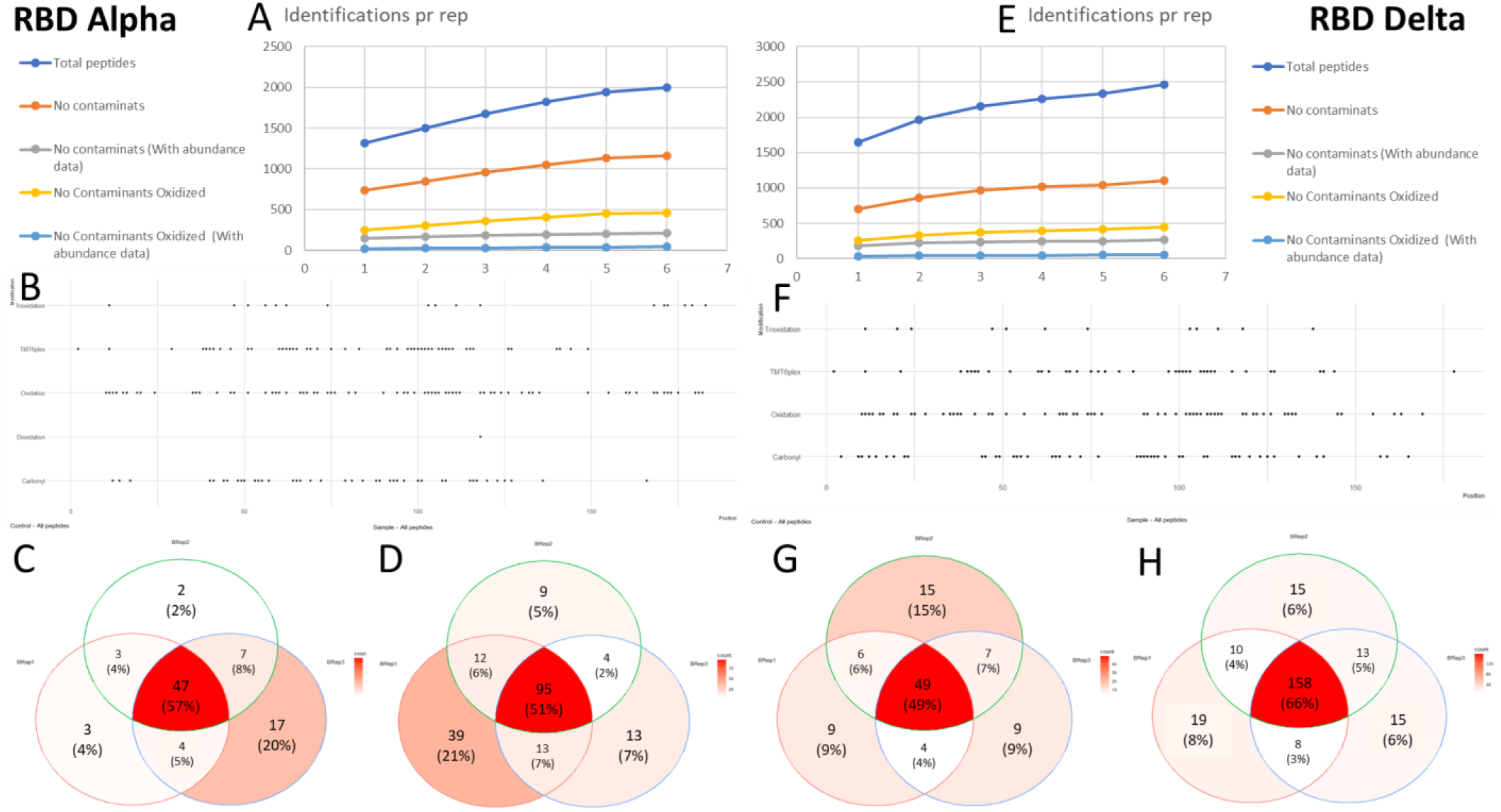
A and E show identified peptide groups depending on the number of technical replicates tested for RBD Alpha and Delta, respectively. Total peptide groups identified (Total peptides), no contaminants (Total, without contaminants), no contaminants with abundance data, no contaminants with oxidations, and no contaminants and oxidized with abundance data. B and F show the distribution of the modifications identified on RBD Alpha (B) and RBD Delta (F), modifications shown are Trioxidation, TMT-labeling, Oxidation, Dioxidation, and Carbonylation. C, D, G, and H show Venn Diagrams of biological replicates, for RBD Alpha (C and D) and RBD Delta (G and H), showing their overlap for Control and Sample experiments.

The modification patterns of RBD Alpha and Delta are shown in Figure 2B and F. Similar patterns were observed, with a high distribution of single oxidations, TMT labelling, and carbonylation. Difference in the tri-oxidation pattern is observed for the last part of the proteins, however, no modifications were observed for RBD Delta from 140 to 190. The TMT labelling was well distributed across the entire protein as expected for satisfactory labeling; however, a few missing areas were observed. The relatively low TMT labelling efficiency of 43% for RBD Alpha and 41% for RBD Delta may have caused the missing areas of labeling. However, this does not affect the conclusions drawn in this paper. The distribution of modified residues for each modification is shown in Supplementary Figure 2. Biological replicates showed a high identification rate of more than 65% in two out of three biological replicates across all samples. The overlap between biological replicates was: for RBD Alpha 70% of control and 67% of sample and for RBD Delta 66% of control and 78% of sample (Figure 2C, D, G, and H).

#### Significant changes

Peptide groups extracted from Proteome Discoverer, with and without oxidative modifications, were analyzed for significant changes between the Ag and Ag + Ab (Mix) samples. All peptide groups identified with significant changes are presented in Table 1. Each peptide group is renamed as peptide and numbered; the complete list is presented in Supplementary Figure 3. The resulting ratios elucidate the changes caused by mixing the two types of antibodies with the antigen. The significance of the changes was tested for each time point (3/0 min and 9/0 min incubation) and the significant peptide ratios are presented in Figure 3. Observed changes are translated into data that either represent a binding or an opening. Binding is achieved by lower abundance in the Ag sample for non-oxidized peptide groups and higher abundance in the Ag sample for oxidized peptide groups compared to the Mix sample. The binding can also be the result of shielding, which cannot be distinguished in this analysis. Binding can be observed between Ag and Ab samples whereas shielding is structural changes within the protein. However, an opening is a region with an increased exposure to the environment e.g., upon binding. An opening is achieved by higher abundance in the Ag sample for non-oxidized peptide groups and lower abundance in the Ag sample for oxidized peptide groups compared to the Mix sample.

**Figure 3.**
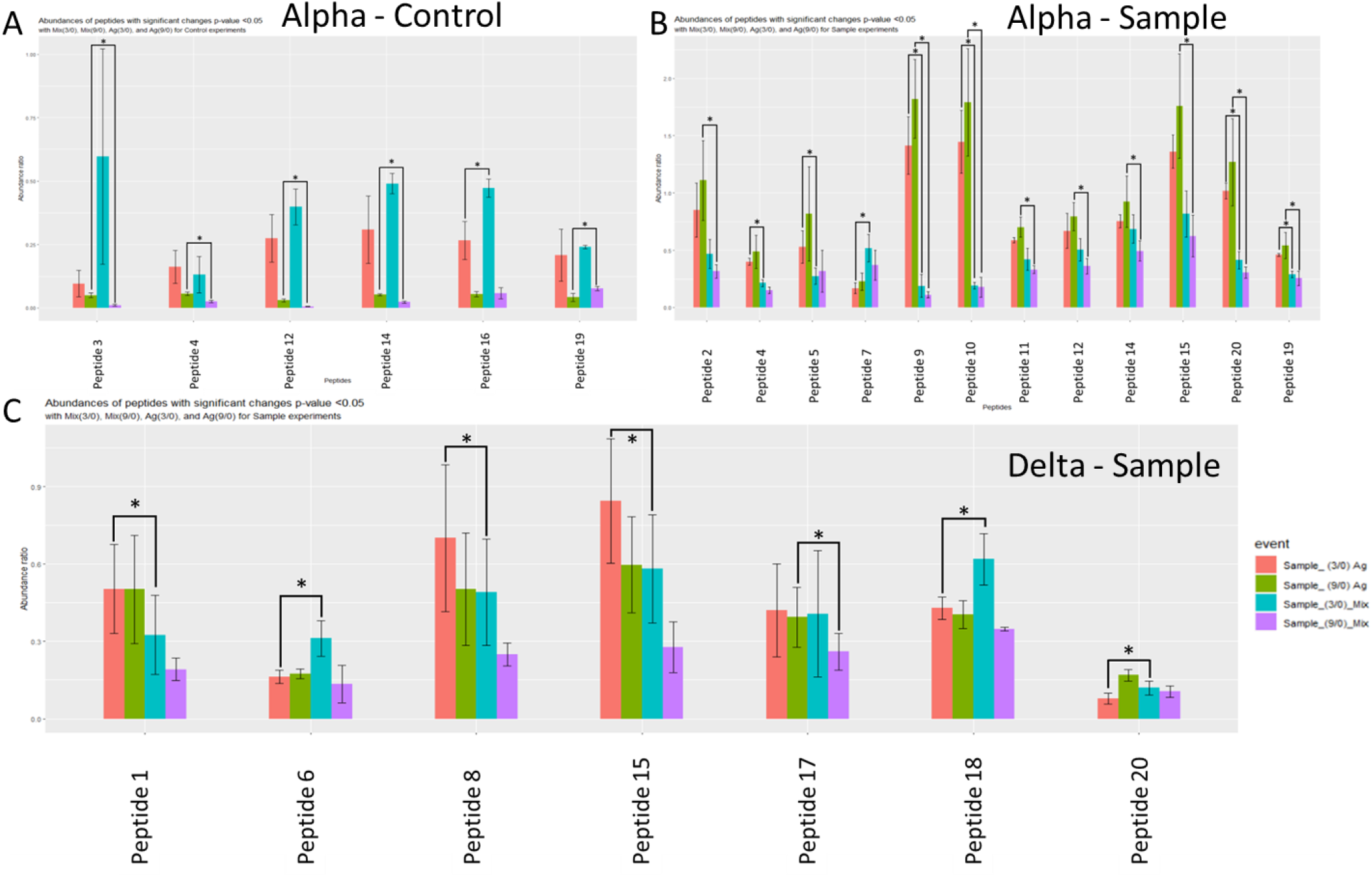
Peptides identified with at least one p-value of less than 0.05 at one time point. The colors represent different time point and sample type: Red = 3/0 min incubation for Ag, Green = 9/0 min incubation for Ag, Cyan = 3/0 min incubation for Mix, Purple = 9/0 min incubation for Mix. A) RBD Alpha Control experiment utilizing negative binding antibodies. B) RBD Alpha Sample experiment utilizing anti-RBD antibodies. C) RBD Delta Sample experiment utilizing anti-RBD antibodies.

**Table 1.** Peptide groups identified for RBD Alpha and Delta Data are concatenated into either showing binding (B) or opening (O). The data shown are from calculating Mix / Ag at 3/0- and 9/0-min incubation. Control and Sample indicates RBD with and without negative control antibody.

| Peptide group name | 1st. residue # | Last residue # | Molecular weight | Modifications | RBD Alpha |  |  |  | RBD Delta |  |  |  | RBD Combined |  |
| --- | --- | --- | --- | --- | --- | --- | --- | --- | --- | --- | --- | --- | --- | --- |
|  |  |  |  |  | Control |  | Sample |  | Control |  | Sample |  | Combined |  |
|  |  |  |  |  | 3/0 | 9/0 | 3/0 | 9/0 | 3/0 | 9/0 | 3/0 | 9/0 | 3/0 | 9/0 |
| Peptide 1 | 328 | 345 | 2328 |  |  |  |  |  |  |  | O |  | O |  |
| Peptide 2 | 356 | 378 | 2758 |  |  |  |  | O |  |  |  |  |  | O |
| Peptide 3 | 357 | 368 | 1632 |  |  | O |  |  |  |  |  |  | - |  |
| Peptide 4 | 357 | 377 | 2602 |  |  | O | O |  |  |  |  |  | - |  |
| Peptide 5 | 357 | 385 | 3495 |  |  |  | O |  |  |  |  |  |  |  |
| Peptide 6 | 378 | 385 | 1140 |  |  |  |  |  |  |  | B |  |  |  |
| Peptide 7 | 392 | 402 | 1513 |  |  |  | B |  |  |  |  |  |  |  |
| Peptide 8 | 396 | 401 | 1036 |  |  |  |  |  |  |  |  | O |  | O |
| Peptide 9 | 396 | 407 | 1592 |  |  |  | O | O |  |  |  |  |  | O |
| Peptide 10 | 400 | 407 | 1172 |  |  |  | O | O |  |  |  |  |  | O |
| Peptide 11 | 408 | 416 | 1128 |  |  | O |  | O |  |  |  |  |  | - |
| Peptide 12 | 424 | 443 | 1227 |  |  |  |  |  |  |  | O |  | - |  |
| Peptide 13 | 424 | 443 | 2495 |  |  | O |  | O |  |  |  |  |  | - |
| Peptide 14 | 424 | 443 | 2527 | 2xOx |  |  |  | B |  |  |  |  |  | B |
| Peptide 15 | 424 | 443 | 2596 |  | B |  |  |  |  |  |  |  | - |  |
| Peptide 16 | 436 | 443 | 1120 |  |  | O |  |  |  |  |  |  |  | - |
| Peptide 17 | 495 | 507 | 1656 |  |  |  |  |  |  |  | B |  | B |  |
| Peptide 18 | 508 | 514 | 1116 |  | B |  | O | O |  |  |  |  | - | O |
| Peptide 19 | 508 | 534 | 3362 |  |  |  | O | O |  |  | B |  | ? | O |

Significant changes were observed for 7 peptides for RBD Alpha Control, with 2 peptides observed at 3/0 min and 5 peptides at 9/0 min incubation. Observed significant peptides span residues 357 to 514 on the intact spike protein. For RBD Alpha Sample, 11 peptides were identified spanning residues 356 to 534. Three peptides were found significantly changed at 3/0 min incubation and 4 peptides were found significantly changed at 9/0 min incubation. The remaining 4 peptides were found at both 3/0 min and 9/0 min incubation. A single peptide (Peptide 15) was identified with double oxidation and was identified at 9/0 min incubation. For RBD Delta, 6 peptides were observed with significant changes at 3/0 min incubation and 1 peptide at 9/0 min incubation, with peptides spanning from residue 328 to 534. Unlike the RBD Alpha Control, the RBD Delta Control did not show any significantly changed peptides.

The data from both RBD Alpha and Delta were combined to establish a common binding site, as we assume that the binding site is the same for both variants. Furthermore, a peptide has been categorized as a false positive if found in the Control experiment for RBD Alpha. Only the RBD Alpha Control experiment has been taken into consideration as no significant peptides were identified for RBD Delta Control. The non-binding negative control antibody used in the control experiments was confirmed to be non-binding by SPR shown in Figure 1. An overview of the peptides identified can be seen in Table 1. Calculations for bindings or openings can be seen in Supplementary Figure 3. The resulting peptides identified with significant changes are peptide 1, 7, 9, 10, and 17 at 3/0 min incubation and peptide 2, 8, 9, 10, 14, 18, and 19 at 9/0 min incubation. Peptides that are found to have opposite data (binding or opening) have been marked with a question mark and are left out of the following analysis. Peptide 7 was identified to be involved in binding but is overlapping with three peptides showing an opening spanning residue 378 to 402. However, the data presented here show only overlapping binding and opening and are therefore only conclusive for residues 392 to 395.

### 3.4 Binding site

Changes found to be significant were mapped to the three-dimensional structure of the spike protein using the pdb structure 7mzg [44][Figure 4]. The peptides identified as significantly changed were mapped to the three-dimensional structure for the RBD Alpha Control experiment [Figure 4A]. Binding was observed at 3/0 min of incubation for two parallel beta sheets, peptide 15 and peptide 18, with unstructured regions in each end. However, several regions show opening for 9/0 min of incubation, including one of the same regions that showed binding but with another peptide, peptide 13. The other regions identified with an opening are peptide 3, 4, 11, and 16. None of the identified peptides were found as significant at both time points.

**Figure 4.**
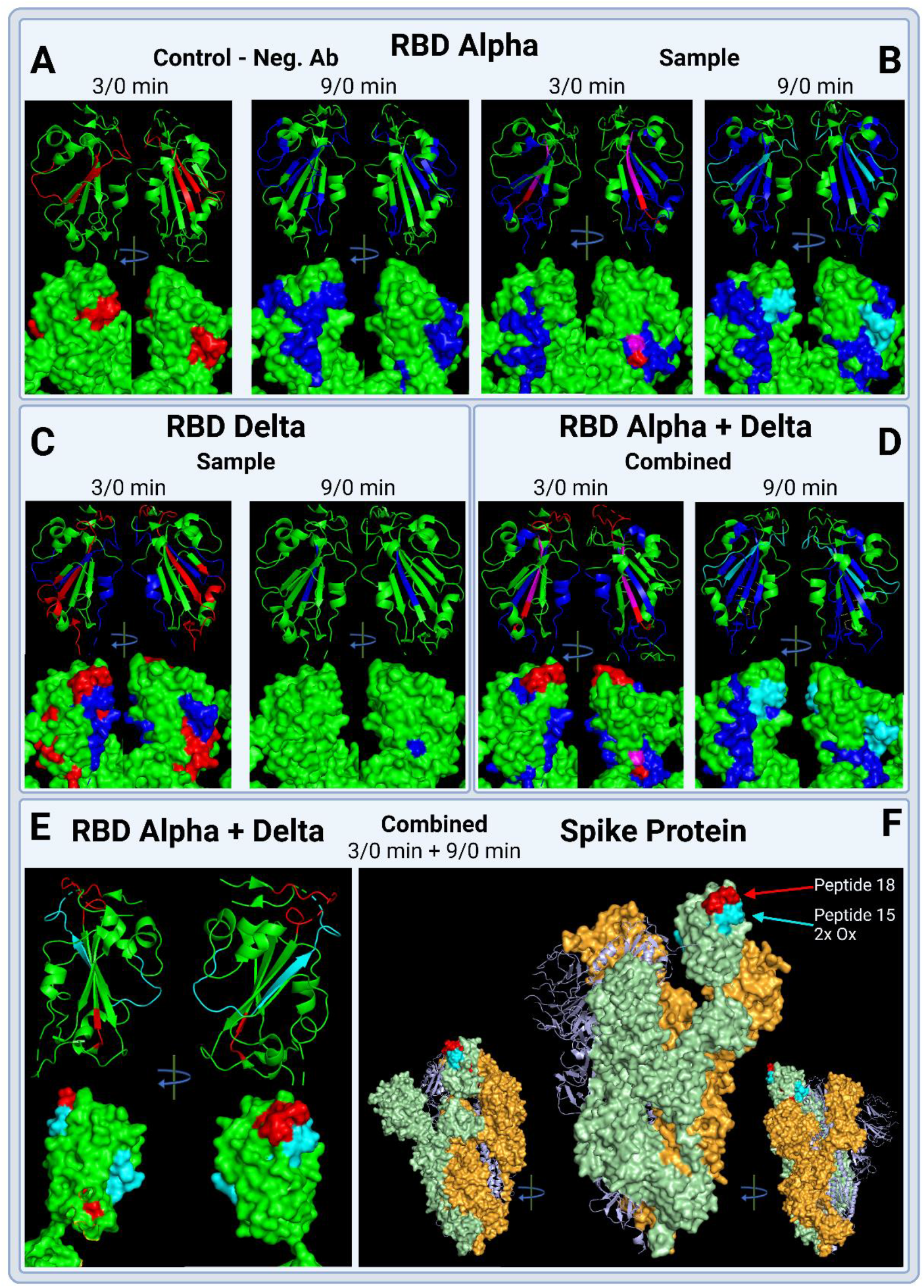
Three-dimensional structure of the Receptor-Binding Domain (RDB) from the SARS-CoV2 spike protein. The pdb structure pdb7mzg was utilized to map the significant changes. The changes are colored the following: Red = Binding/shielding (Decreased exposure), Blue = Opening (Increased exposure), Magenta = found to be both binding and opening, Cyan = Binding of double oxidized peptide. A) RBD Alpha Control experiment with negative antibodies for 3/0 min and 9/0 min incubation. B) RBD Alpha Sample experiment with anti-RBD antibodies for 3/0 min and 9/0 min incubation. C) RBD Delta Sample experiment with anti-RBD antibodies. C) Combined data for the RBD Alpha and RBD Delta Sample experiment with anti-RBD antibodies for 3/0 min and 9/0 min incubation. Data from the Control experiment have been subtracted from the combined data. E) Combined data for RBD Alpha and RBD Delta Sample experiment with anti-RBD antibodies for 3/0 min and 9/0 min incubation. Only the binding data are shown. F) Combined data for the RBD Alpha and RBD Delta Sample experiment with anti-RBD antibodies for 3/0 min and 9/0 min incubation. Mapped to the complete spike protein. Each protein was distinguished by color; green, orange, and light purple, respectively.

For the RBD Alpha Sample experiment, Figure 4B, the regions of residue 396 to 407 and 508 to 534 (peptides 9 and 10) were found to show opening upon binding to the antibody for both time points, 3/0 min and 9/0 min incubation. Openings were also observed for region 357 to 385 at 3/0 min incubation (peptide 4 and 5), whereas, for 9/0 min incubation openings were observed for regions 356 to 378, 400 to 416, and 424 to 443 (peptide 2, 11, and 13). Binding was observed for the Alpha Sample experiment at region 392 to 402 for 3/0 min incubation (peptide 7), and 424 to 443 for 9/0 min incubation when a double oxidation is present (peptide 14).

For the RBD Delta Sample experiment, Figure 4C, openings were observed for regions 328 to 345 and 424 to 443 for 3/0 min incubation (peptide 1 and 12). Binding at 3/0 min incubation was observed for region 378 to 385 and 495 to 534 (peptide 6, 17, and 19). For 9/0 min incubation, only region 436 to 443 was observed with an opening (peptide 8).

The significant changes observed for RBD Alpha and Delta were combined as described previously and mapped to the three-dimensional structure shown in Figure 4D. This data show two primary binding sites, one in the top of the RBD, generally accessible, and the second located on the inside towards the center of the trimer. To simplify the binding site, only the regions showing binding were mapped to the structure and the two time points were combined, shown in Figure 4E. Binding of anti-RBD to RBD Alpha and Delta was mapped to region 495 to 507 and the double oxidized region 424 to 443, peptide 17 and 14, respectively. The binding was additionally mapped to the trimer of the spike protein, where it was mapped to the open conformation of the spike protein, Figure 4F.

## 4 Discussion

The currently used methods within HRPF analysis of protein-protein interactions have a few shortcomings. The problems in particular are linked to a wide variety of modification outcomes, low dynamic range, and reliability of the quantification used. We therefore set out to make an experimental setup of epitope mapping with hydroxyl radical protein footprinting (HRPF) using Fenton chemistry, that could be reliable while still using relative long incubation times (3 min and 9 min). The following controls were introduced to increase the reliability and robustness of the data: no oxidation controls (0 min incubation) for all samples, six times technical replicates, biological triplicates, and a negative control antibody that does not bind the antigen. Unlike the majority of current HRPF experimental setups [23–27] we introduced a stable isotope labelling strategy by incorporating TMT labelling for relative quantification.

Quantification of HRPF data is problematic due to the known introduction of many PTMs across the entire protein, thereby increasing the complexity of the sample. Furthermore, the stoichiometry of the introduced oxidation and other PTMs are low, which is problematic when using data dependent acquisition (DDA) methods. The current DDA method incorporated a long exclusion time in addition to 6 technical replicates for all samples in order to increase the depth of the analysis. Compared to the literature, most studies using FPOP focus on a single non-oxidation control and biological triplicates [35,45]. Alternatively, biological duplicates for both oxidized and non-oxidized control have been used [23,46]. New technologies have recently been introduced, employing HRPF for epitope mapping. These methods have generally used 3 biological replicates for both oxidized and non-oxidized samples [29,30]. Common to all the above-mentioned works is the analysis of two proteins and their binding epitope, where each protein is analysed alone and in combination. However, none of these studies included negative non-binding control antibodies (neg. Ab) or decoy proteins. The use of these neg. Ab or decoy proteins could prove essential in minimalizing the false discovery rate. We have shown in Figure 4A that significant changes can be recorded even though no binding was observed by SPR, Figure 1. The use of neg. Ab is common practice within the antibody research field, as antibodies are notorious for having unspecific binding. We therefore encourage others to utilize this in their work, as it proved to exclude several peptides from the final analysis e.g., peptide 3, 4, 11, 12, 13, 15, and 16. This reduced the significantly changed peptides from 19 to 12, corresponding to a false positive rate of 37% of the identified peptides with significant changes.

The TMT labelling strategy allowed us to decrease the number of samples to be processed and at the same time limit the number of missing values across samples. Furthermore, using a TMT common channel we were able to normalize all samples and directly compare ratios across sample types and between control and sample experiments. The relative quantification showed that both increases and decreases in abundance could be analysed for both non-oxidized and oxidized peptides, corresponding to binding and openings of regions.

The present epitope mapping approach is based on Fenton chemistry and is not to the authors’ knowledge commercially available through contract research organizations. Other methodologies using hydroxyl radical protein footprinting (HRPF) are available both within academia and commercially. Fast Photochemical Oxidation of Proteins (FPOP), invented by Gross’ laboratory [25,28], promised a wide use, but due to high cost of implementation and safety hazards of the laser the method has not been put into practice in the standard laboratory [29]. Therefore, several other applications for HRPF have been suggested with X-ray activated HRPF by Jain et al. [27], Flash Oxidation system (FOX) by Sharp et al. [29], and Plasma-Induced Modification of Biomolecules (PLIMB) by Sussman et al. [30]. All the methods mentioned above have the same shortcoming regarding implementation; a specific instrument is needed to perform the chemistry prior to sample preparation and analysis by mass spectrometry. For the development of our approach we set out to make the experimental setup more robust and integrate commonly used standards from both antibody research and proteomics with regards to negative control antibodies, biological and technical replicates.

The main drawback of Fenton chemistry, which has been addressed several times in the available literature regarding HRPF, is the time scale of the reaction, ranging from several seconds to minutes [29]. This time scale does not allow for investigation of protein dynamics but could prove useful when identifying antibody-antigen interactions due to their high stability. The introduction of controls on several levels combined with semi-automatic sample handling by a pipetting robot, achieves a high level of reproducibility and robustness. The method presented in the present work should be widely applicable and easily implementable as the only additional equipment needed is a pipetting robot. Such equipment has recently become more common due to increased sample load combined with faster mass spectrometers.

The binding epitope identified by our experimental setup found that anti-RBD bound to both RBD Alpha and Delta at residues 392 to 402, 424 to 443 when double oxidized, and 495 to 507, corresponding to peptide 7, 14, and 17, respectively. The region identified here as the binding epitope matches that found by both Yuan et al. [47] and Wu et al. [48]. The binding site mapped to the complete trimer of the SARS-CoV-2 spike protein, shown in Figure 4F, showed that the binding site is located on the tip of the receptor-binding domain and is accessible in both closed and open structure [49].

## 5 Conclusion

While the use of hydroxyl radical protein footprinting (HRPF) is a relatively commonly used method for determining protein-protein interactions, the inclusion of a negative protein control has not been widely applied. However, the use of nonsense or negative antibody control is routinely used in immune-chemistry and antibody research. In the current project we demonstrated that by introducing negative antibody controls, 37% of the identified peptides with a significant change were false positives when using HRPF method for epitope mapping. Direct comparison across sample types and conditions were applicable due to TMT labeling. Furthermore, the introduction of TMT labeling, semi-automatic sample handling, biological triplicates, and six technical replicates, achieved data with great depth of analysis and reliability.

The binding epitope identified on the receptor-binding domain from the SARS-CoV-2 spike protein variant Alpha and Delta, was residue 392 to 402, 424 to 443, and 495 to 507, represented by peptides 7, 14, and 17, respectively. This binding site matches the one found in the previous epitope mapping studies of anti-RBD SARS-CoV-2 neutralizing antibodies. A potential binding site, residue 508 to 514, was ruled out due to the same region being positive in the negative antibody control.

In conclusion we suggest that the combination of experimental controls and stable isotope labelling strategy could be implemented widely to ensure the limitation of false discovery in the HRPF epitope mapping. This will considerably minimize the risk of wrongly assigned peptides in binding sites and therefore be of great use and inspiration to increase the reliability of mass spectrometric based epitope mapping methodologies.

## Supplementary Data

### A.1

**A)**

HHHHHHHHHHFTVEKGIYQTSNFRVQPTESIVRFPNITNLCPFGEVFNATRFASVYAWNRKRISNCVADYSVLYNSASFSTFKCYGVSPTKLNDLCFTNVYADSFVIRGDEVRQIAPGQTGKIADYNYKLPDDFTGCVIAWNSNNLDSKVGGNYNYLYRLFRKSNLKPFERDISTEIYQAGSTPCNGVEGFNCYFPLQSYGFQPTNGVGYQPYRVVVLSFELLHAPATVCGPKKSTNLVKNKCVNFNF

Coverage 92.74%; total spectra 166; unique spectra 89.

**B)**

HHHHHHHHHHFTVEKGIYQTSNFRVQPTESIVRFPNITNLCPFGEVFNATRFASVYAWNRKRISNCVADYSVLYNSASFSTFKCYGVSPTKLNDLCFTNVYADSFVIRGDEVRQIAPGQTGKIADYNYKLPDDFTGCVIAWNSNNLDSKVGGNYNYLYRLFRKSNLKPFERDISTEIYQAGSTPCNGVEGFNCYFPLQSYGFQPTNGVGYQPYRVVVLSFELLHAPATVCGPKKSTNLVKNKCVNFNF

Coverage 100%; total spectra 196; unique spectra 95.

**C)**

MGILPSPGMPALLSLVSLLSVLLMGVAFRVQPTESIVRFPNITNLCPFGEVFNATRFASVYAWNRKRISNCVADYSVLYNSASFSTFKCYGVSPTKLNDLCFTNVYADSFVIRGDEVRQIAPGQTGKIADYNYKLPDDFTGCVIAWNSNNLDSKVGGNYNYRYRLFRKSNLKPFERDISTEIKQAGSTPCNGVEGFNCYFPLQSYGFQPTNGVGYQPYRVVVLSFELLHAPATVCGPKKSTNLVKNKCVNFGHHHHHHH

Coverage 76.83%; total spectra 1532; unique spectra 2.

*Supplementary Figure 1. Sequence coverage obtained for various receptor-binding domains of SARS-CoV-2 spike variants by tryptic digests. A) Native SARS-CoV-2 Spike protein Wild Type (Wuhan). B) PNGase F treated SARS-CoV-2 Spike protein Wild Type (Wuhan). The deamidated asparagines in N(IT) and N(AT) show that these residues are glycosylated in the native protein. C) SARS-CoV-2 Spike protein Delta. Green shows identified residues, black unidentified residues, and red deamidated asparagine residues*.

### A.2

**Supplementary Figure 1.**
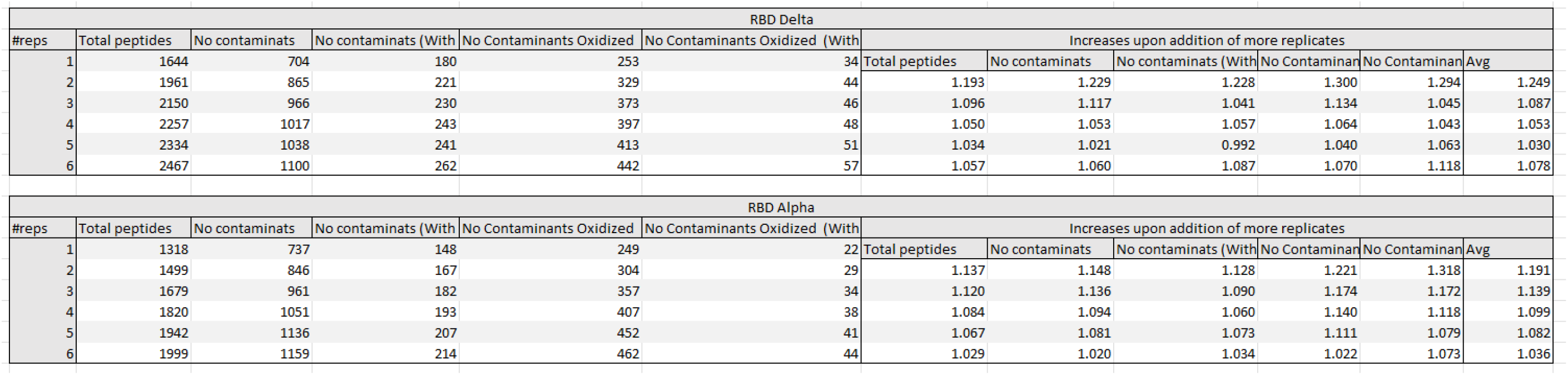
Identified peptide groups in different subcategories for each additional technical replicate. Increases of each technical replicate can be seen in the last column below “ Increases upon addition of more replicates”.

### A.3

**Supplementary Figure 2.**
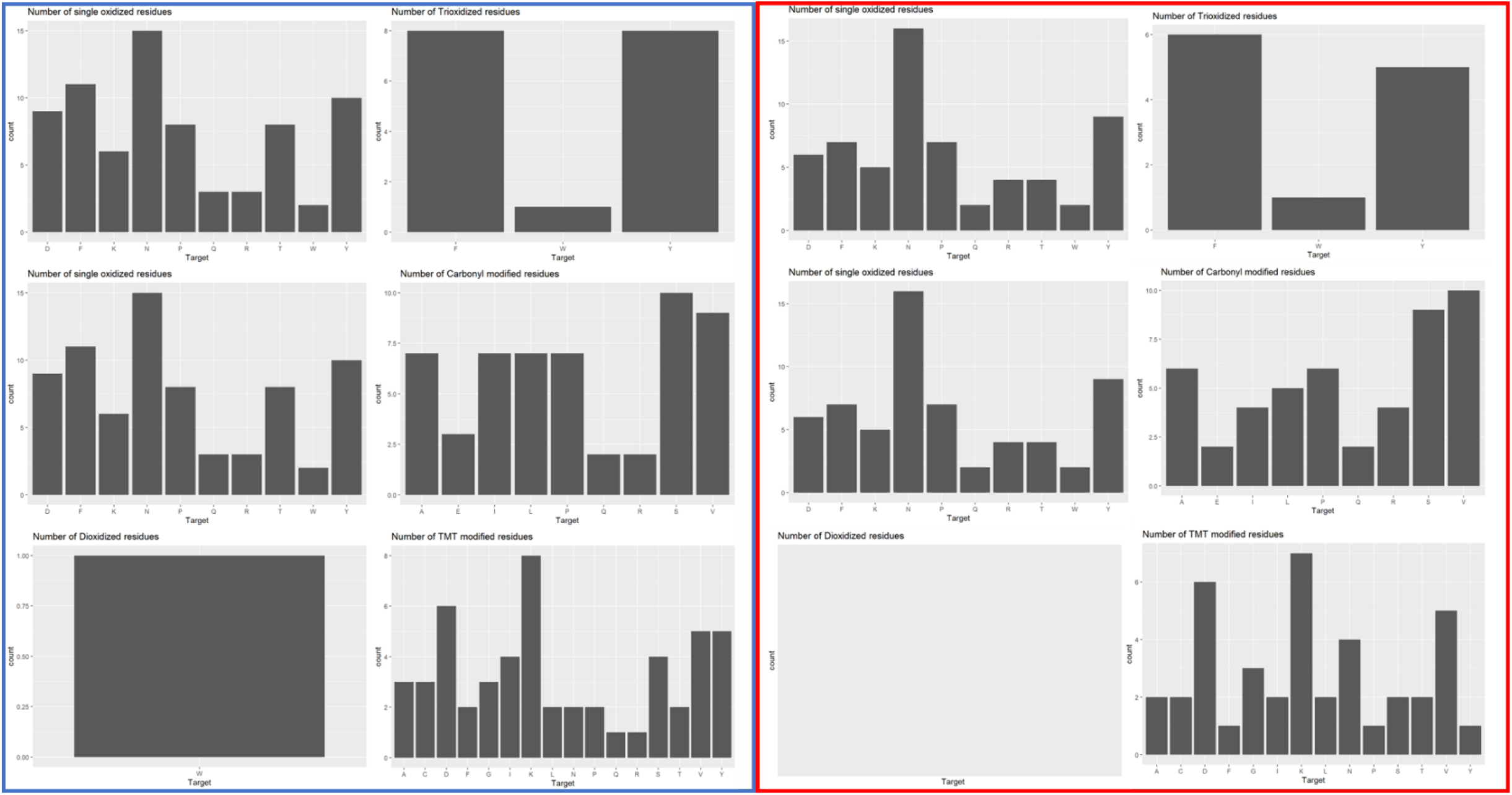
Identified modifications and their residue distribution for each RBD Wuhan (Blue) and RBD Delta (Red). Modifications shown are: Mono oxidation, di-oxidation, tri-oxidation, carbonylation, and TMT labeling.

### A.4

**Supplementary Figure 3.**
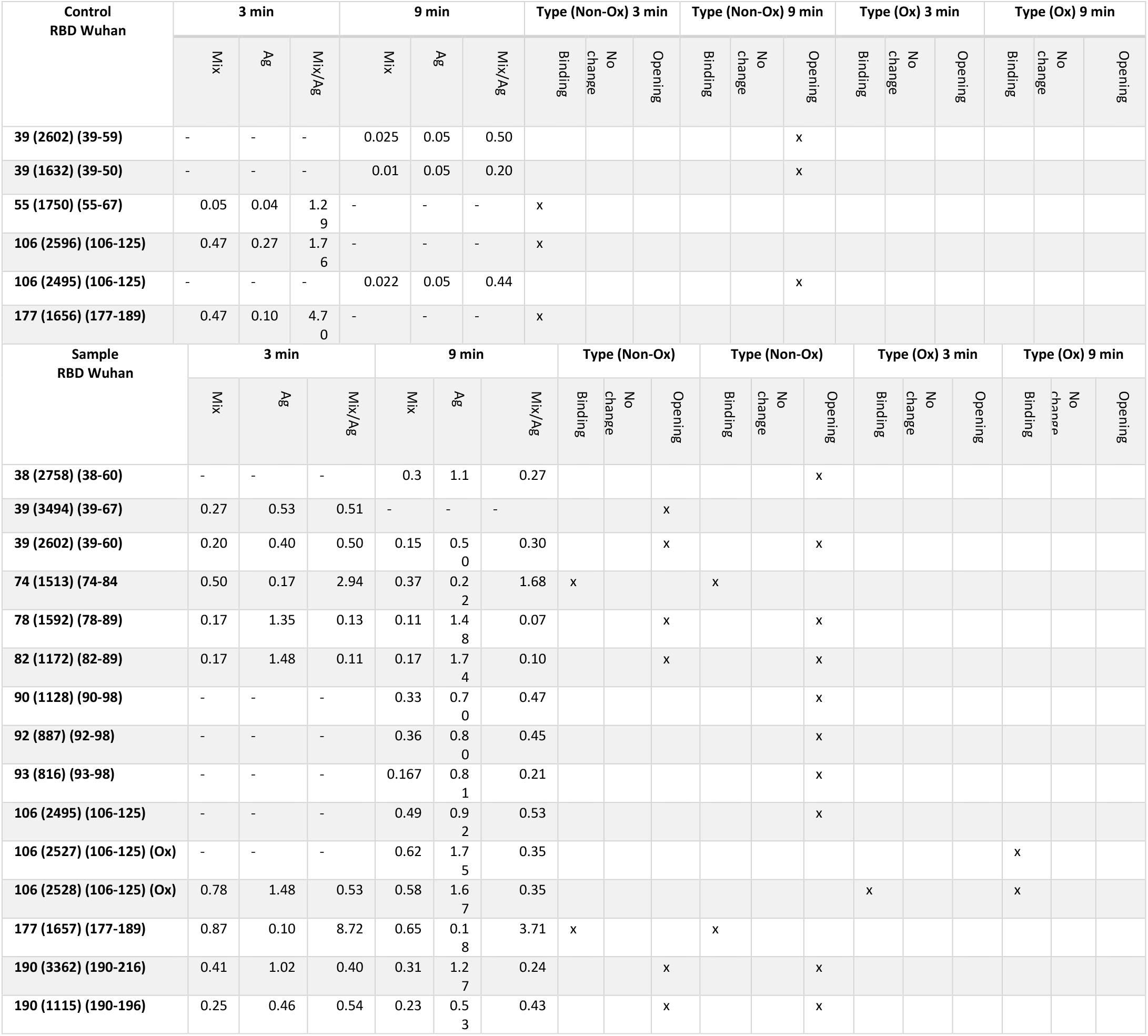
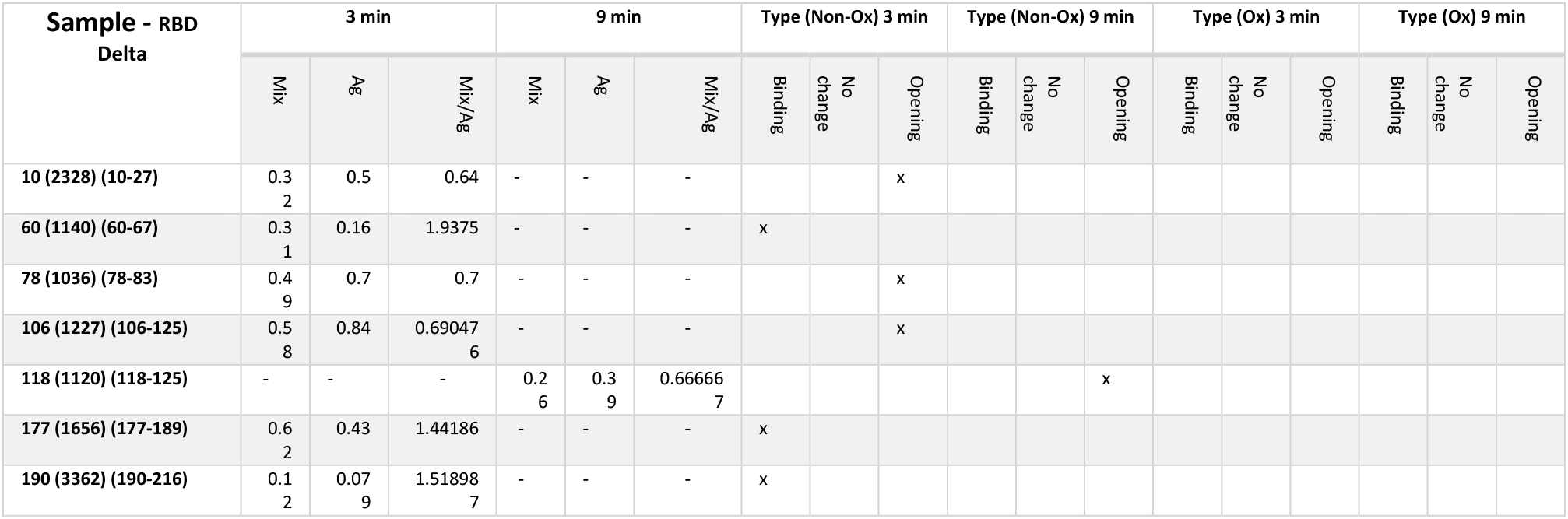
Table of identified peptides with a p-value < 0.05 and their corresponding ratio abundances. Mig/Ag abundances are calculated, and type of binding/shielding, no change, or an opening is marked. Data for both Control and Sample experiments for RBD Wuhan is displayed along with data from Sample experiment for RBD Delta, as no significant changes were observed for RBD Delta Control experiment.

## References

[1] A.E. Gorbalenya, S.C. Baker, R.S. Baric, R.J. de Groot, C. Drosten, A.A. Gulyaeva, B.L. Haagmans, C. Lauber, A.M. Leontovich, B.W. Neuman, D. Penzar, S. Perlman, L.L.M. Poon, D. V. Samborskiy, I.A. Sidorov, I. Sola, J. Ziebuhr, The species Severe acute respiratory syndrome-related coronavirus: classifying 2019-nCoV and naming it SARS-CoV-2, Nat. Microbiol., 5 (2020) 536–544.

[2] D. Wrapp, N. Wang, K.S. Corbett, J.A. Goldsmith, C.-L. Hsieh, O. Abiona, B.S. Graham, J.S. McLellan, Cryo-EM structure of the 2019-nCoV spike in the prefusion conformation, Science (80-.)., 367 (2020) 1260–1263.

[3] V. Monteil, H. Kwon, P. Prado, A. Hagelkrüys, R.A. Wimmer, M. Stahl, A. Leopoldi, E. Garreta, C. Hurtado del Pozo, F. Prosper, J.P. Romero, G. Wirnsberger, H. Zhang, A.S. Slutsky, R. Conder, N. Montserrat, A. Mirazimi, J.M. Penninger, Inhibition of SARS-CoV-2 Infections in Engineered Human Tissues Using Clinical-Grade Soluble Human ACE2, Cell, 181 (2020) 905–913.e7.

[4] R. Shi, C. Shan, X. Duan, Z. Chen, P. Liu, J. Song, T. Song, X. Bi, C. Han, L. Wu, G. Gao, X. Hu, Y. Zhang, Z. Tong, W. Huang, W.J. Liu, G. Wu, B. Zhang, L. Wang, J. Qi, et al., A human neutralizing antibody targets the receptor-binding site of SARS-CoV-2, Nature, 584 (2020) 120–124.

[5] C.G. Rappazzo, L. V. Tse, C.I. Kaku, D. Wrapp, M. Sakharkar, D. Huang, L.M. Deveau, T.J. Yockachonis, A.S. Herbert, M.B. Battles, C.M. O’Brien, M.E. Brown, J.C. Geoghegan, J. Belk, L. Peng, L. Yang, Y. Hou, T.D. Scobey, D.R. Burton, D. Nemazee, et al., Broad and potent activity against SARS-like viruses by an engineered human monoclonal antibody, Science (80-.)., 371 (2021) 823–829.

[6] D. Pinto, Y.-J. Park, M. Beltramello, A.C. Walls, M.A. Tortorici, S. Bianchi, S. Jaconi, K. Culap, F. Zatta, A. De Marco, A. Peter, B. Guarino, R. Spreafico, E. Cameroni, J.B. Case, R.E. Chen, C. Havenar-Daughton, G. Snell, A. Telenti, H.W. Virgin, et al., Cross-neutralization of SARS-CoV-2 by a human monoclonal SARS-CoV antibody, Nature, 583 (2020) 290–295.

[7] F. Bertoglio, D. Meier, N. Langreder, S. Steinke, U. Rand, L. Simonelli, P.A. Heine, R. Ballmann, K.-T. Schneider, K.D.R. Roth, M. Ruschig, P. Riese, K. Eschke, Y. Kim, D. Schäckermann, M. Pedotti, P. Kuhn, S. Zock-Emmenthal, J. Wöhrle, N. Kilb, et al., SARS-CoV-2 neutralizing human recombinant antibodies selected from pre-pandemic healthy donors binding at RBD-ACE2 interface, Nat. Commun., 12 (2021) 1577.

[8] I.A. Favorskaya, D. V. Shcheblyakov, I.B. Esmagambetov, I. V. Dolzhikova, I.A. Alekseeva, A.I. Korobkova, D. V. Voronina, E.I. Ryabova, A.A. Derkaev, A. V. Kovyrshina, A.A. Iliukhina, A.G. Botikov, O.L. Voronina, D.A. Egorova, O. V. Zubkova, N.N. Ryzhova, E.I. Aksenova, M.S. Kunda, D.Y. Logunov, B.S. Naroditsky, et al., Single-Domain Antibodies Efficiently Neutralize SARS-CoV-2 Variants of Concern, Front. Immunol., 13 (2022).

[9] Food and Drug Administration Fact sheet for healthcare providers: Emergency Use Authorization for Evusheld (tixagevimab co-packaged with cilgavimab, (n.d.).

[10] Food and Drug Administration Fact sheet for healthcare providers: Emergency Use Authorization (EUA) of sotrovimab, FDA, (n.d.).

[11] I. Petta, S. Lievens, C. Libert, J. Tavernier, K. De Bosscher, Modulation of Protein–Protein Interactions for the Development of Novel Therapeutics, Mol. Ther., 24 (2016) 707–718.

[12] N.T. Ditto, B.D. Brooks, The emerging role of biosensor-based epitope binning and mapping in antibody-based drug discovery, Expert Opin. Drug Discov., 11 (2016) 925–937.

[13] S. Hayes, B. Malacrida, M. Kiely, P.A. Kiely, Studying protein–protein interactions: progress, pitfalls and solutions, Biochem. Soc. Trans., 44 (2016) 994–1004.

[14] M. Zhou, Q. Li, R. Wang, Current Experimental Methods for Characterizing Protein–Protein Interactions, ChemMedChem, 11 (2016) 738–756.

[15] Z. Zhang, D.L. Smith, Determination of amide hydrogen exchange by mass spectrometry: A new tool for protein structure elucidation, Protein Sci., 2 (1993) 522–531.

[16] V. Katta, B.T. Chait, S. Carr, Conformational changes in proteins probed by hydrogen-exchange electrospray-ionization mass spectrometry, Rapid Commun. Mass Spectrom., 5 (1991) 214–217.

[17] R.S. Johnson, K.A. Walsh, Mass spectrometric measurement of protein amide hydrogen exchange rates of apo- and holo-myoglobin, Protein Sci., 3 (1994) 2411–2418.

[18] Y. Kostyukevich, T. Acter, A. Zherebker, A. Ahmed, S. Kim, E. Nikolaev, Hydrogen/deuterium exchange in mass spectrometry, Mass Spectrom. Rev., 37 (2018) 811–853.

[19] F.J. O’Reilly, J. Rappsilber, Cross-linking mass spectrometry: methods and applications in structural, molecular and systems biology, Nat. Struct. Mol. Biol., 25 (2018) 1000–1008.

[20] M. Schneider, A. Belsom, J. Rappsilber, Protein Tertiary Structure by Crosslinking/Mass Spectrometry, Trends Biochem. Sci., 43 (2018) 157–169.

[21] Z.A. Chen, J. Rappsilber, Protein structure dynamics by crosslinking mass spectrometry, Curr. Opin. Struct. Biol., 80 (2023).

[22] C. Yu, L. Huang, Cross-Linking Mass Spectrometry: An Emerging Technology for Interactomics and Structural Biology, Anal. Chem., 90 (2018) 144–165.

[23] D.S. Loginov, J. Fiala, P. Brechlin, G. Kruppa, P. Novak, Hydroxyl radical footprinting analysis of a human haptoglobin-hemoglobin complex, Biochim. Biophys. Acta - Proteins Proteomics, 1870 (2022).

[24] M.L. Gau, Brian and Garai, Kanchan and Frieden, Carl and Gross, Mass spectrometry-based protein footprinting characterizes the structures of oligomeric apolipoprotein E2, E3, and E4, Biochemistry, 50 (2011) 8117–8126.

[25] M.L. Li, Ke Sherry and Shi, Liuqing and Gross, Mass spectrometry-based fast photochemical oxidation of proteins (FPOP) for higher order structure characterization, Acc. Chem. Res., 51 (2011) 736–744.

[26] J. Kiselar, M.R. Chance, High-Resolution Hydroxyl Radical Protein Footprinting: Biophysics Tool for Drug Discovery, (2018).

[27] R. Jain, N.S. Dhillon, E.R. Farquhar, B. Wang, X. Li, J. Kiselar, M.R. Chance, Multiplex Chemical Labeling of Amino Acids for Protein Footprinting Structure Assessment, Anal. Chem., 94 (2022) 9819–9825.

[28] B. Gau, K. Garai, C. Frieden, M.L. Gross, Mass spectrometry-based protein footprinting characterizes the structures of oligomeric apolipoprotein E2, E3, and E4, Biochemistry, 50 (2011) 8117–8126.

[29] J.S. Sharp, E.E. Chea, S.K. Misra, R. Orlando, M. Popov, R.W. Egan, D. Holman, S.R. Weinberger, Flash Oxidation (FOX) System: A Novel Laser-Free Fast Photochemical Oxidation Protein Footprinting Platform, J. Am. Soc. Mass Spectrom., 32 (2021) 1601–1609.

[30] B.B. Minkoff, J.M. Blatz, F.A. Choudhury, D. Benjamin, J.L. Shohet, M.R. Sussman, Plasma-Generated OH Radical Production for Analyzing Three-Dimensional Structure in Protein Therapeutics, Sci. Rep., 7 (2017).

[31] H.J.H. Fenton, On a new reaction of tartaric acid, Chem. News, 33 (1876) 190.

[32] W.H. Koppenol, The Haber-Weiss cycle - 70 years later, Redox Rep., 6 (2001) 229–234.

[33] C.Y. Ralston, J.S. Sharp, Structural Investigation of Therapeutic Antibodies Using Hydroxyl Radical Protein Footprinting Methods, Antibodies, 11 (2022).

[34] M. Lin, D. Krawitz, M.D. Callahan, G. Deperalta, A.T. Wecksler, Characterization of ELISA Antibody-Antigen Interaction using Footprinting-Mass Spectrometry and Negative Staining Transmission Electron Microscopy, J. Am. Soc. Mass Spectrom., 29 (2018) 961–971.

[35] A.J. Schick, V. Lundin, J. Low, K. Peng, R. Vandlen, A.T. Wecksler, Epitope mapping of anti-drug antibodies to a clinical candidate bispecific antibody, MAbs, 14 (2022).

[36] N. Shembekar, H. Hu, D. Eustace, C.A. Merten, Single-Cell Droplet Microfluidic Screening for Antibodies Specifically Binding to Target Cells, Cell Rep., 22 (2018) 2206–2215.

[37] T. Pochechueva, A. Chinarev, N. Bovin, A. Fedier, F. Jacob, V. Heinzelmann-Schwarz, PEGylation of microbead surfaces reduces unspecific antibody binding in glycan-based suspension array, J. Immunol. Methods, 412 (2014) 42–52.

[38] M. Polák, G. Yassaghi, D. Kavan, F. Filandr, J. Fiala, Z. Kukačka, P. Halada, D.S. Loginov, P. Novák, Utilization of Fast Photochemical Oxidation of Proteins and Both Bottom-up and Top-down Mass Spectrometry for Structural Characterization of a Transcription Factor-dsDNA Complex, Anal. Chem., 94 (2022) 3203–3210.

[39] R.G.H. Cotton, C. Milstein, Fusion of Two Immunoglobulin-producing Myeloma Cells, Nature, 244 (1973) 42–43.

[40] S. Heissel, S.J. Frederiksen, J.B. Id, H. Peter, Enhanced trypsin on a budget: Stabilization, purification and high-temperature application of inexpensive commercial trypsin for proteomics applications, (2019) 1–16.

[41] J. Rappsilber, M. Mann, Y. Ishihama, Protocol for micro-purification, enrichment, pre-fractionation and storage of peptides for proteomics using StageTips, Nat. Protoc., 2 (2007) 1896.

[42] M. Leser, J.R. Chapman, M. Khine, J. Pegan, M. Law, M. El Makkaoui, B.M. Ueberheide, M. Brenowitz, Chemical Generation of Hydroxyl Radical for Oxidative ‘Footprinting,’ Protein Pept. Lett., 26 (2019) 61–69.

[43] Y. Perez-Riverol, J. Bai, C. Bandla, D. García-Seisdedos, S. Hewapathirana, S. Kamatchinathan, D.J. Kundu, A. Prakash, A. Frericks-Zipper, M. Eisenacher, M. Walzer, S. Wang, A. Brazma, J.A. Vizcaíno, The PRIDE database resources in 2022: a hub for mass spectrometry-based proteomics evidences, Nucleic Acids Res., 50 (2022) D543–D552.

[44] A.K. Wheatley, P. Pymm, R. Esterbauer, M.H. Dietrich, W.S. Lee, D. Drew, H.G. Kelly, L.-J. Chan, F.L. Mordant, K.A. Black, A. Adair, H.-X. Tan, J.A. Juno, K.M. Wragg, T. Amarasena, E. Lopez, K.J. Selva, E.R. Haycroft, J.P. Cooney, H. Venugopal, et al., Landscape of human antibody recognition of the SARS-CoV-2 receptor binding domain, Cell Rep., 37 (2021) 109822.

[45] M. Lin, D. Krawitz, M.D. Callahan, G. Deperalta, A.T. Wecksler, Characterization of ELISA Antibody-Antigen Interaction using Footprinting-Mass Spectrometry and Negative Staining Transmission Electron Microscopy, J. Am. Soc. Mass Spectrom., 29 (2018) 961–971.

[46] R. Jiang, D.L. Rempel, M.L. Gross, MALDI peptide mapping for fast analysis in protein footprinting, Int. J. Mass Spectrom., 490 (2023).

[47] M. Yuan, H. Liu, N.C. Wu, C.-C.D. Lee, X. Zhu, F. Zhao, D. Huang, W. Yu, Y. Hua, H. Tien, T.F. Rogers, E. Landais, D. Sok, J.G. Jardine, D.R. Burton, I.A. Wilson, Structural basis of a shared antibody response to SARS-CoV-2, Science (80-.)., 369 (2020) 1119–1123.

[48] Y. Wu, F. Wang, C. Shen, W. Peng, D. Li, C. Zhao, Z. Li, S. Li, Y. Bi, Y. Yang, Y. Gong, H. Xiao, Z. Fan, S. Tan, G. Wu, W. Tan, X. Lu, C. Fan, Q. Wang, Y. Liu, et al., A noncompeting pair of human neutralizing antibodies block COVID-19 virus binding to its receptor ACE2, Science (80-.)., 368 (2020) 1274–1278.

[49] I.M. Francino-Urdaniz, T.A. Whitehead, An overview of methods for the structural and functional mapping of epitopes recognized by anti-SARS-CoV-2 antibodies, RSC Chem. Biol., 2 (2021) 1580–1589.

